# *ATP-dependent Clp protease subunit C1, HvClpC1*, is a strong candidate gene for barley variegation mutant *luteostrians* as revealed by genetic mapping and genomic re-sequencing

**DOI:** 10.1101/2021.02.04.429718

**Authors:** Mingjiu Li, Ganggang Guo, Hélène Pidon, Michael Melzer, Alberto R. Prina, Thomas Börner, Nils Stein

**Author notes:** **Corresponding authors**: Mingjiu Li; Nils Stein.

## Abstract

Implementation of next-generation sequencing in forward genetic screens greatly accelerated gene discovery in species with larger genomes, including many crop plants. In barley, extensive mutant collections are available, however, the causative mutations for many of the genes remains largely unknown. Here we demonstrate how a combination of low-resolution genetic mapping, whole-genome resequencing and comparative functional analyses provides a promising path towards candidate identification of genes involved in plastid biology and / or photosynthesis, even if genes are located in recombination poor regions of the genome. As a proof of concept, we simulated the prediction of a candidate gene for the recently cloned variegation mutant *albostrians* (*HvAST* / *HvCMF7*) and adopted the approach for suggesting *HvClpC1* as candidate gene for the yellow-green variegation mutant *luteostrians*.

**Author Summary:** Forward genetics is an approach of identifying a causal gene for a mutant phenotype and has proven to be a powerful tool for dissecting the genetic control of biological processes in many species. A large number of barley mutants was generated in the 1940s to 1970s when mutation breeding programs flourished. Genetic dissection of the causative mutations responsible for the phenotype, however, lagged far behind, limited by lack of molecular markers and high-throughput genotyping platforms. Next-generation sequencing technologies have revolutionized genomics, facilitating the process of identifying mutations underlying a phenotype of interest. Multiple mapping-by-sequencing or cloning-by-sequencing strategies were established towards fast gene discovery. In this study, we used mapping-by-sequencing to identify candidate genes within coarsely delimited genetic intervals, for two variegation mutants in barley – *luteostrians* and *albostrians*. After testing the approach using the example of the previously cloned *albostrians* gene *HvAST*, the gene *HvClpC1* could be delimited as candidate gene for *luteostrians.* The mapping-by-sequencing strategy implemented here is generally suited for surveying barley mutant collections for phenotypes affecting fundamental processes of plant morphology, physiology and development.

## Introduction

Barley mutagenesis was intensely studied in the mid twentieth century. These activities resulted in extensive mutant collections available through genebanks such as NordGen (https://www.nordgen.org/) and IPK (https://gbis.ipk-gatersleben.de/gbis2i/faces/index.jsf). Barley mutants served as valuable resource for dissecting the genetic basis of a wide range of complex biological processes. Their broad use, however, was impeded until recently by a lack of genomic resources and tools. This has been changed by the fast development of next-generation sequencing (NGS) based technologies and strategies for gene cloning. Due to diminishing sequencing costs, NGS can be applied for sequencing-based genotyping of whole mapping populations as well as for whole-genome resequencing to accelerate gene discovery even in large genome crop species (Candela et al., 2015; Jaganathan et al., 2020). Mapping-by-sequencing strategies initially were not always ready applicable to all species as they could be limited. For instance, the Mutmap (Abe et al., 2012) and Mutmap+ (Fekih et al., 2013) approaches, initially applied to rice, require access to a high-quality reference genome for the genotype used for mutagenesis; the homozygosity mapping approach requires availability of genotypes with a known pedigree (Singh et al., 2013). Special attention needs to be paid to experimental design and technical decisions in order to enforce that sequencing data will allow to map a mutation of interest (Wilson-Sanchez et al., 2019). In barley, *MANY-NODED DWARF* (*MND*) was cloned by a technique similar to SHOREmap (Schneeberger et al., 2009); *LAXATUM-a* (*LAX-a*) was isolated by exome-capture sequencing (Mascher et al., 2013) of several highly informative recombinant pools (Jost et al., 2016). Notably, identification of *MND* and *LAX-a* was even achieved while relying on a largely unordered draft genome of barley (International Barley Genome Sequencing Consortium, 2012). Recently, the release and improvement of a high-quality reference genome of barley (Mascher et al., 2017; Monat et al., 2019) has greatly facilitated forward genetic screens relying largely on mapping-/ cloning-by-sequencing strategies (Candela et al., 2015) and thus paved the way towards systematic dissection of the genic factors underlying barley mutant resources.

Many mutants of genebank collections are affected by photosynthesis-related defects leading to aberrant coloration phenotypes. Photosynthesis-related mutants provide a highly valuable genetic tool for identification of nuclear genes involved in different aspects of chloroplast development. Among the distinct chlorophyll-deficient phenotypes, leaf variegation is a common phenomenon that has been observed for many plants in nature (Toshoji et al., 2012). Investigations with variegated mutants (patched in dicots / striping in monocots) generated insights into the molecular mechanisms of leaf variegation (Yu et al., 2007). A threshold-dependent genetic model was proposed as a mechanism underlying *var2* variegation. *Var2* encodes a chloroplast-localized protein AtFtsH2 which belongs to the *filamentation temperature sensitive* (*FtsH*) *metalloprotease* gene family (Chen et al., 2000; Takechi et al., 2000). In the model, two pairs of FtsH proteins, AtFtsH1/5 and AtFtsH2/8, form oligomeric complexes in the thylakoid membrane and a threshold level of oligomeric complexes is required for normal chloroplast function and green sector formation (Yu et al., 2004). In monocot species, *iojap* of maize (*Zea mays*) (Walbot and Coe, 1979) and *albostrians* of barley (Hagemann and Scholz, 1962; Hess et al., 1993) represent two examples of classical variegation mutation revealing that cells of albino sectors contain ribosome-free plastids. *iojap* encodes a component associated with the plastid ribosomal 50S subunit (Han et al., 1992). The *albostrians* gene *HvAST* / *HvCMF7* encodes a plastid-localized CCT MOTIF FAMILY (CMF) protein (Li et al., 2019c). Although a threshold-dependent mechanism is also very likely, the molecular mechanisms underlying the *iojap* and *albostrians* leaf variegation still remain elusive. *luteostrians* is another barley mutant with a block in early chloroplast development, showing a yellow-green striped variegation phenotype that appears only if the mutant allele is inherited through the female gamete.

Here we report *HvClpC1* as a candidate gene for the variegation mutant *luteostrians* by applying a sequencing-based gene identification strategy that has great potential of systematical application to reveal causative mutations for photosynthesis-related mutants in barley collections.

## Results

### Inheritance of Leaf Variegation in the *luteostrians* Mutant

Phenotypically, progeny of *luteostrians* can be classified into three categories: 1) green-yellow striped, 2) completely yellow, and 3) albino (Figure 1A). The green-yellow striped seedlings were able to complete the life cycle while the yellow and albino plants do not survive beyond the three-leaf vegetative growth stage. Considering the variegated pattern to be analogous to the *albostrians* mutant of barley (Li et al., 2019c), we named the causal gene underlying the *luteostrians* striped phenotype as *HvLST*, representing *Hordeum vulgare LUTEOSTRIANS*. Selfing of heterozygous plants leads to about a quarter of aborted grains, indicating that homozygosity for the gene (*lst/lst*) is lethal at post-zygote or early embryonic stage (Table 1). Inheritance of the yellow striped phenotype was gamete-dependent (Figure 1B) since variegation is observed if the *lst* allele is inherited through the female gamete (i.e., *lst/LST*). This interpretation is supported by three lines of evidence: 1) when wild type (*LST/LST*) plants are fertilized with pollen from striped (*lst/LST*) or green heterozygous plants (*LST/lst*), in either case, only half of the F1 plants offspring showed phenotypic segregation in F2 generation; 2) a segregation ratio of 2:1 (green:striped) was observed in segregant offspring in F2 generation; and 3) heterozygous F2 plants (*LST/lst* or *lst/LST*) segregated with green (*LST/LST* or *LST/lst*) and striped (*lst/LST*) progenies with a ratio of 2:1 in subsequent F3 generation (Figure 1B). Thus, in addition to its essential function during embryogenesis, we postulate that *HvLST* plays an essential role in plastid differentiation / programming during gametic stage in the egg cell.

**Figure 1.**
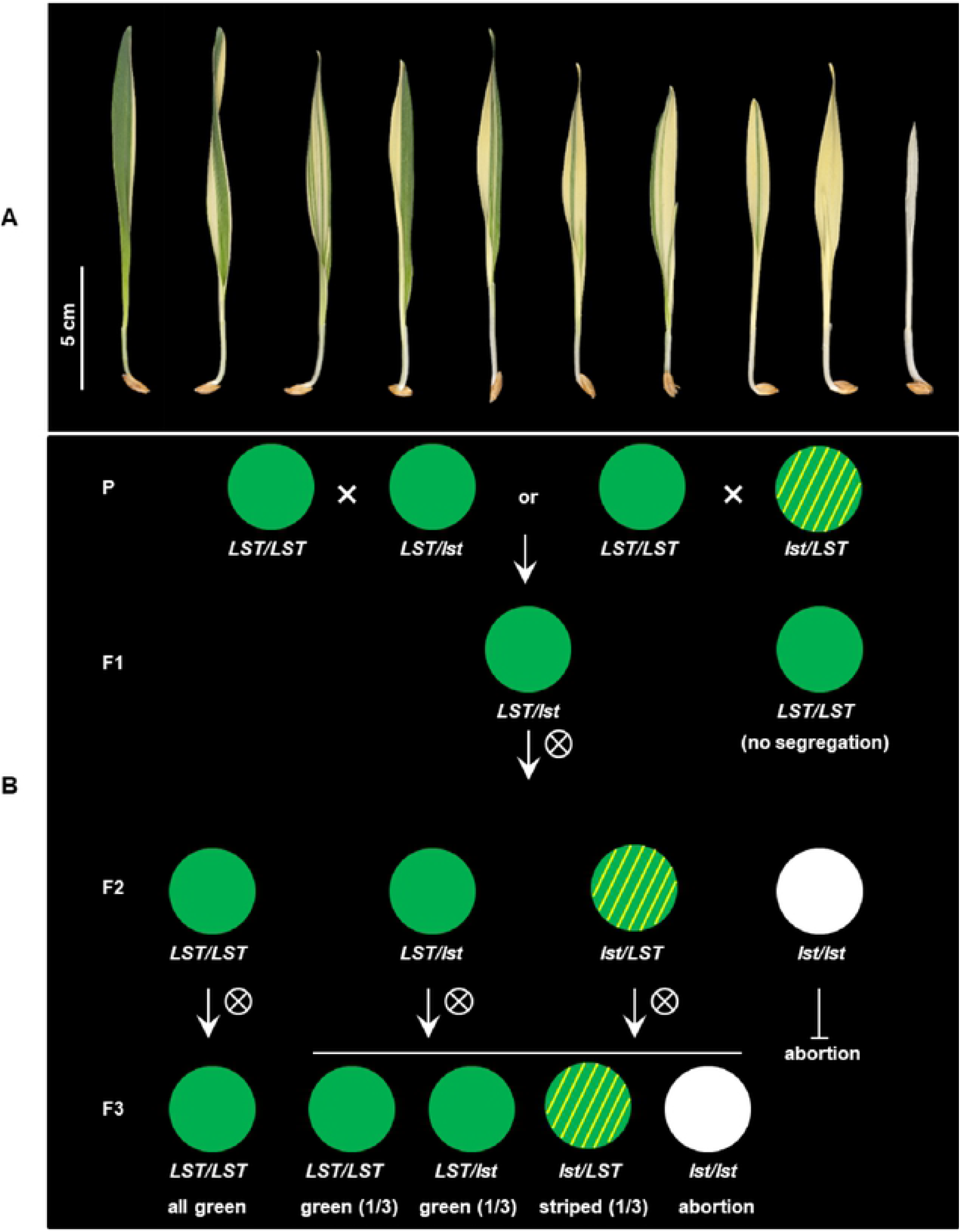
Phenotype and inheritance of variegation in the barley mutant *luteostrians*. (A) Penetration of the mutant phenotype varies among seedlings, ranging from a narrow yellow stripe to complete yellowish or albino. Neither yellowish/albino plants survived beyond third-leaf stage. (B) Inheritance pattern of the *luteostrians* mutant phenotype. Variegation only occurs in plants if the *lst* allele was transmitted through the female gamete. Upper panel: Heterozygous plants can be obtained by using either green or variegated plants (heterozygous for the *luteostrians* allele) as pollen donor. This will generate 50% F1-progeny heterozygous for *luteostrians* (panel F1). Progenies of selfed F1 heterozygotes will exhibit Mendelian segregation in F2; the variegated phenotype, however, will appear only in 50% of the heterozygous plants, carrying the mutant allele inherited from F1 female gamete. Zygotes homozygous for the *luteostrians* allele will be aborted as homozygosity of *luteostrians* early zygotic lethal. Lower panel: Green phenotype of homozygous wild type plants in F2 will be stably transmitted in F3; progenies of heterozygous F2 plants follow a Mendelian inheritance pattern in F3.

**Table 1.**
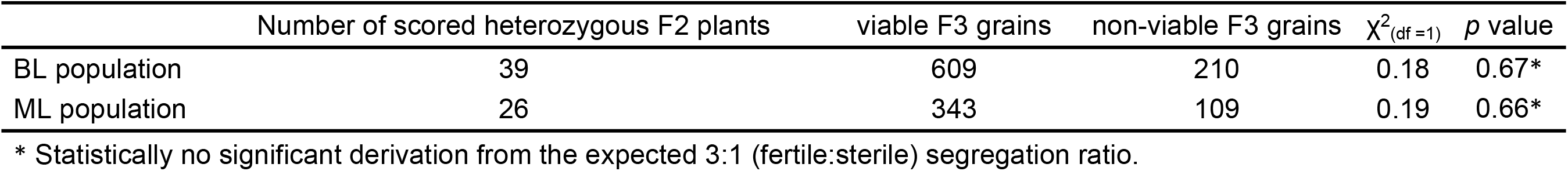
Phenotypic segregation of zygotic lethality in F2 of BL and ML populations.

| | Number of scored heterozygous F2 plants | viable F3 grains | non-viable F3 grains | $\chi^2_{(df=1)}$ | *p* value |
| --- | --- | --- | --- | --- | --- |
| BL population | 39 | 609 | 210 | 0.18 | 0.67* |
| ML population | 26 | 343 | 109 | 0.19 | 0.66* |

\* Statistically no significant derivation from the expected 3:1 (fertile:sterile) segregation ratio.

### Chloroplast Translation is Abolished in the *luteostrians* Mutant

Defective chloroplasts do not contain 70S ribosomes in the *albostrians* mutant (Hess et al., 1993; Li et al., 2019c). In an attempt to check the function of the translation machinery in plastids of the *luteostrians* mutant, we initially examined accumulation of the rRNAs in wild type and *luteostrians* mutant. In analogy to the *albostrians* mutant, the 16S and 23S rRNA species are not observed in defective plastids of the *luteostrians* mutant (Figure 2), indicating the lack of 70S ribosomes and consequently missing chloroplast translation in plastids of the yellow leaf sectors. Next, we studied chloroplast/plastid ultrastructure by transmission electron microscopy. These analyses revealed normal chloroplast development in wild type and in green leaf sections of the *luteostrians* mutant; in both cases chloroplasts contained well-developed stroma and grana thylakoids (Figure 3A & 3B). In contrast, plastids in yellow sectors of the variegated *luteostrians* leaves contained no grana and only rudimentary stroma lamellae (Figure 3C). Notably, chloroplast 70S ribosomes were not detectable in yellow plastids of the *luteostrians* mutant (Figure 3). Altogether, based on the absence of chloroplast rRNA species and the lack of 70S ribosomes it can be postulated that chloroplast translation is abolished in the *luteostrians* mutant.

**Figure 2.**
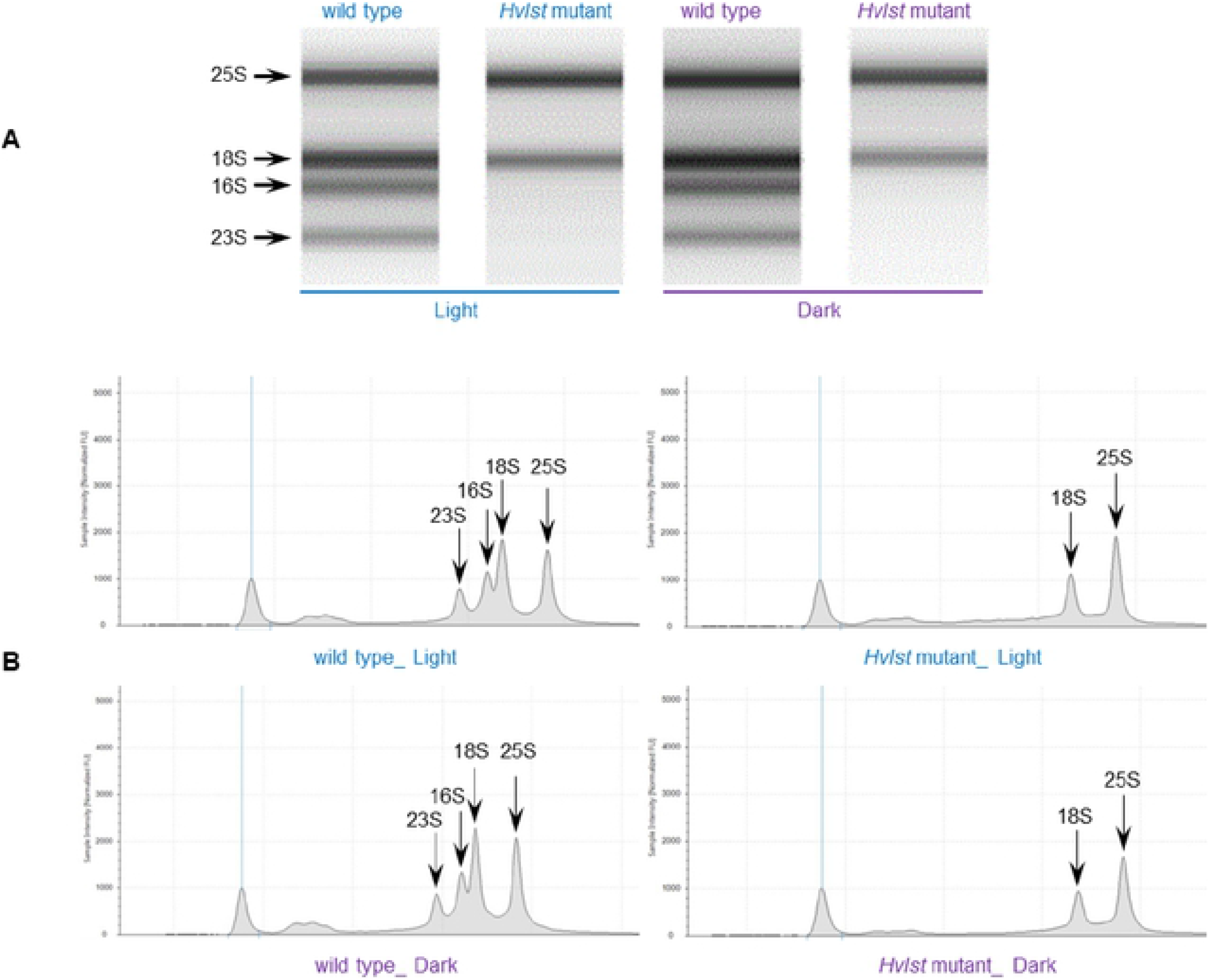
rRNA analysis of wild type and Hvlst mutant. (A) Separation of cytosolic and plastid rRNAs using the Agilent high sensitivity RNA ScreenTape assay. (B) Analysis of rRNA from wild type and *Hvlst* mutant using Agilent Tapestation 4200. The respective cytosolic and plastid rRNA species are indicated by arrows.

**Figure 3.**
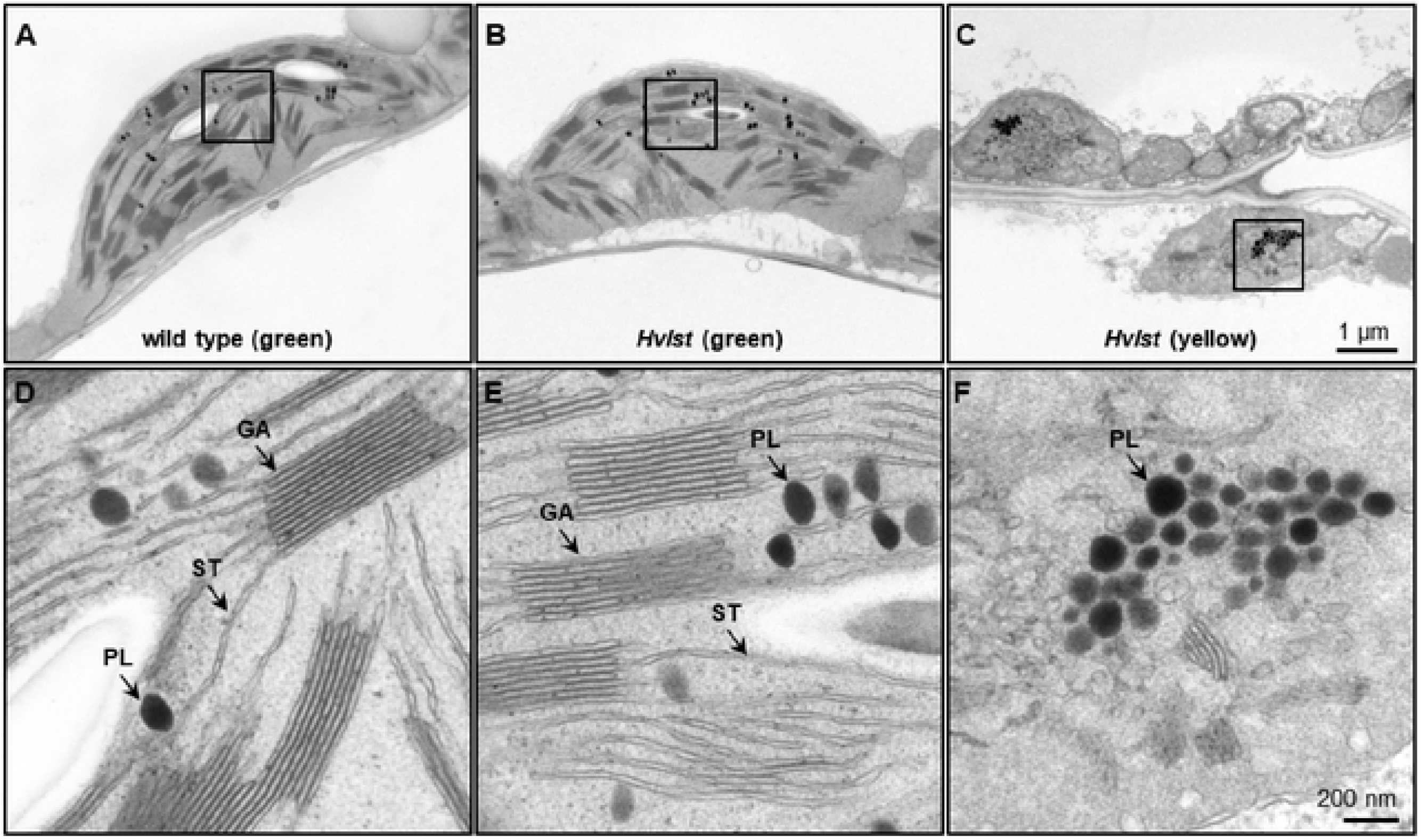
Ultrastructural analysis of chloroplasts of the wild type and Hvlst mutant. (A-C) Ultrastructural analysis of wild type (A), and green (B) and yellow (C) sectors of the *Hvlst* mutant, respectively. Wild type and green leaves of the *Hvlst* mutant contain chloroplasts with fully differentiated grana and stroma thylakoids. By contrast, in plastids of yellow leaves of the *Hvlst* mutant only some pieces of undeveloped membranes were observed. (D-F) Larger magnification of square areas of the corresponding plastid in the top panels A to C. GA, grana; ST, stroma thylakoid; PL, plastoglobuli.

### Genetic Mapping of *HvLST* to the Genetic Centromere of Chromosome 2H

Two F2 mapping populations, designated as ‘BL’ and ‘ML’, were constructed for the purpose of genetic mapping of the *HvLST* gene. The genotypic status of the *luteostrians* locus of F2 plants was determined by phenotypic segregation analysis of their respective F3 progenies. As homozygosity (*lst/lst*) leads to grain abortion, the BL population contained 95 wild type (*LST/LST*) and 172 heterozygotes (*LST/lst* and *lst/LST*), consistent with the segregation of a single recessive gene (Table 2). Genetic mapping in the ML population was affected by segregation distortion with genotype ratios deviating from the expected Mendelian ratios (Table 2). Genotyping-by-sequencing was performed for 267 and 269 F2 genotypes in BL and ML populations, respectively. Sequencing data was mapped to the reference genome of barley (Monat et al., 2019) for SNP calling. In total, 3,745 and 5,507 SNP markers were obtained genome-wide at a minimum sequencing coverage of six-fold for BL and ML populations, respectively. By applying a permissive threshold of 5% missing data for both molecular marker and F2 genotype, mapping could be performed in 124 F2 / 3369 SNPs and 146 F2 / 4854 SNPs for the BL and ML populations, respectively (Supplemental Table 1). Genetic maps for seven linkage groups, representing the seven barley chromosomes, were obtained for BL (LOD ≥ 6) and ML (LOD = 10) populations, comprising mapped markers for 66.4% (2238/3369) and 71.5% (3469/4584) of the originally defined SNPs, respectively (Supplemental Table 1). The accuracy of the linkage maps was consistency-checked by aligning genetic marker positions with their respective physical order in the reference genome of barley (Monat et al., 2019). The *HvLST* gene was assigned to the region of the genetic centromere region of chromosome 2H with a physical distance between the flanking markers of 461.7 and 499.9 Mbp in BL and ML populations, respectively (Supplemental Figure 1).

**Table 2.** Genotypic segregation of the *HvLST* locus in F2 of BL and ML populations.

| | Maternal | Paternal | Population size | Wild type | Heterozygote | $\chi^2_{(df=1)}$ | *p* value |
| --- | --- | --- | --- | --- | --- | --- | --- |
| BL Population | Barke | *luteostrians (lst/LST)* | 269 | 95 | 172 | 0.61 | 0.44* |
| ML Population | Morex | *luteostrians (lst/LST)* | 271 | 117 | 152 | 12.47 | 0.0004<sup>§</sup> |

\* Statistically no significant derivation from the expected 1:2 (wild type:heterozygote) segregation ratio.
<sup>§</sup> Segregation distortion observed for ML population.

Next, PCR-based KASP (Kompetitive Allele-Specific PCR) markers were designed and employed to saturate the identified *HvLST* intervals in both mapping populations. Initially, genotyping all the 267 (BL population) and 269 (ML population) F2 individuals with two population-specific KASP markers (chr2H_178489464 and chr2H_461744325 for BL population; chr2H_200083049 and chr2H_463274491 for ML population) allowed to confirm and further narrow down the *HvLST* target region. Notably, the numerical value within each marker designation indicates the physical coordinate of the mapped SNP position on the reference genome of barley (Monat et al., 2019) (Figure 4). Subsequently, saturation mapping of the *HvLST* interval was performed in a total of 18 recombinants (13 in BL / 5 in ML) with additional KASP markers that were suitable for both mapping populations (Supplemental Table 2). Finally, we delimited the *HvLST* target region to a 0.74 centiMorgan (cM) interval between flanking markers chr2H_431057673 and chr2H_458001177, spanning a distance of ~26.94 Mbp (Figure 5 and Supplemental Table 2). A cluster of markers co-segregated with the *HvLST* locus, suggesting that maximum genetic resolution was achieved at the given size of both mapping populations (Figure 4).

**Figure 4.**
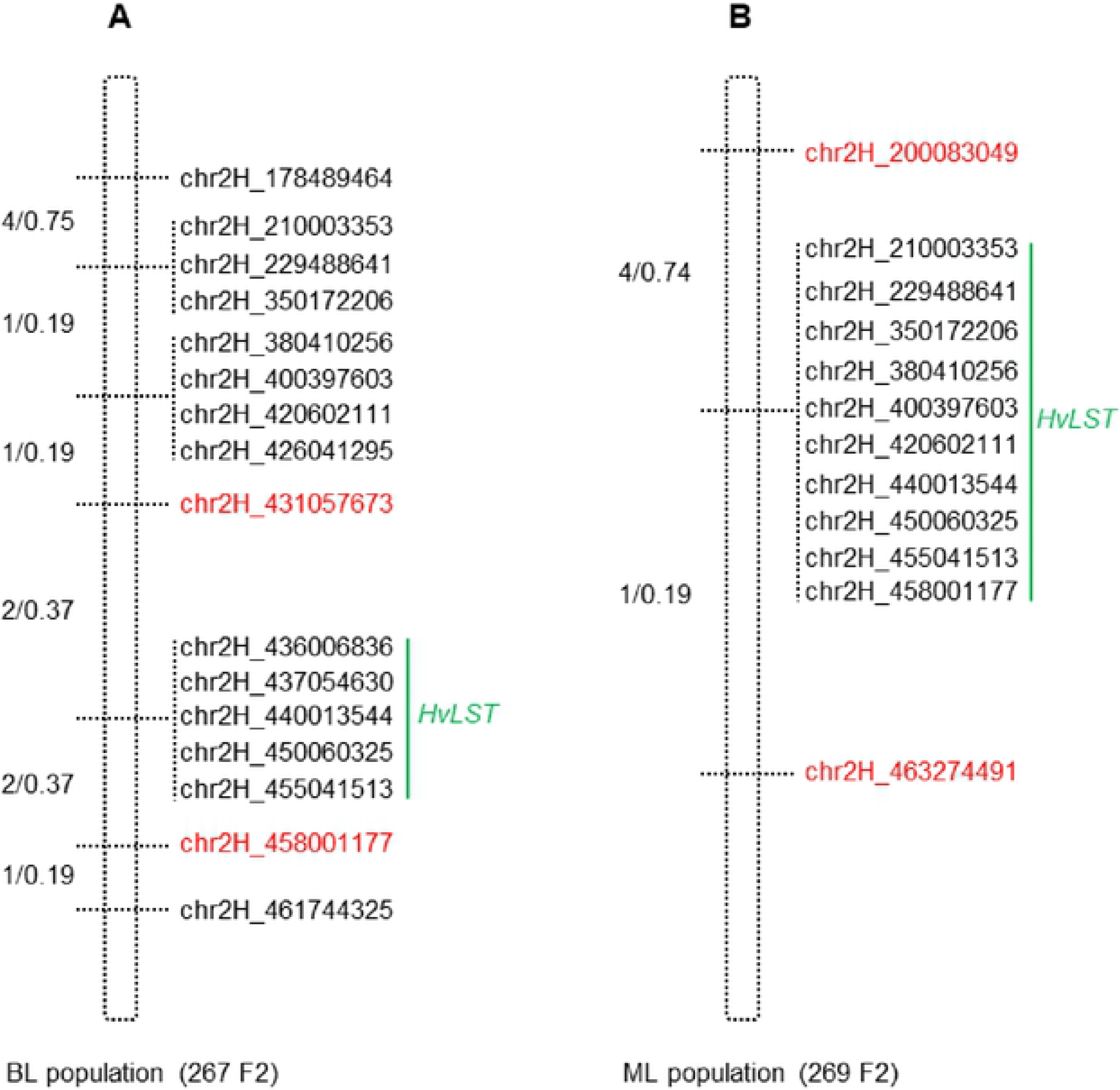
Saturation mapping of the delineated HvLST genetic interval. (A) Marker saturation around the *HvLST* locus in BL population. (B) Marker saturation around the *HvLST* locus in ML population. Recombination events and genetic distance (recombinants/cM) between the neighboring markers are shown on the left of each genetic map. Markers co-segregating with the gene *HvLST* are indicated by a vertical green line. Red font is highlighting the closest flanking markers.

**Figure 5.**
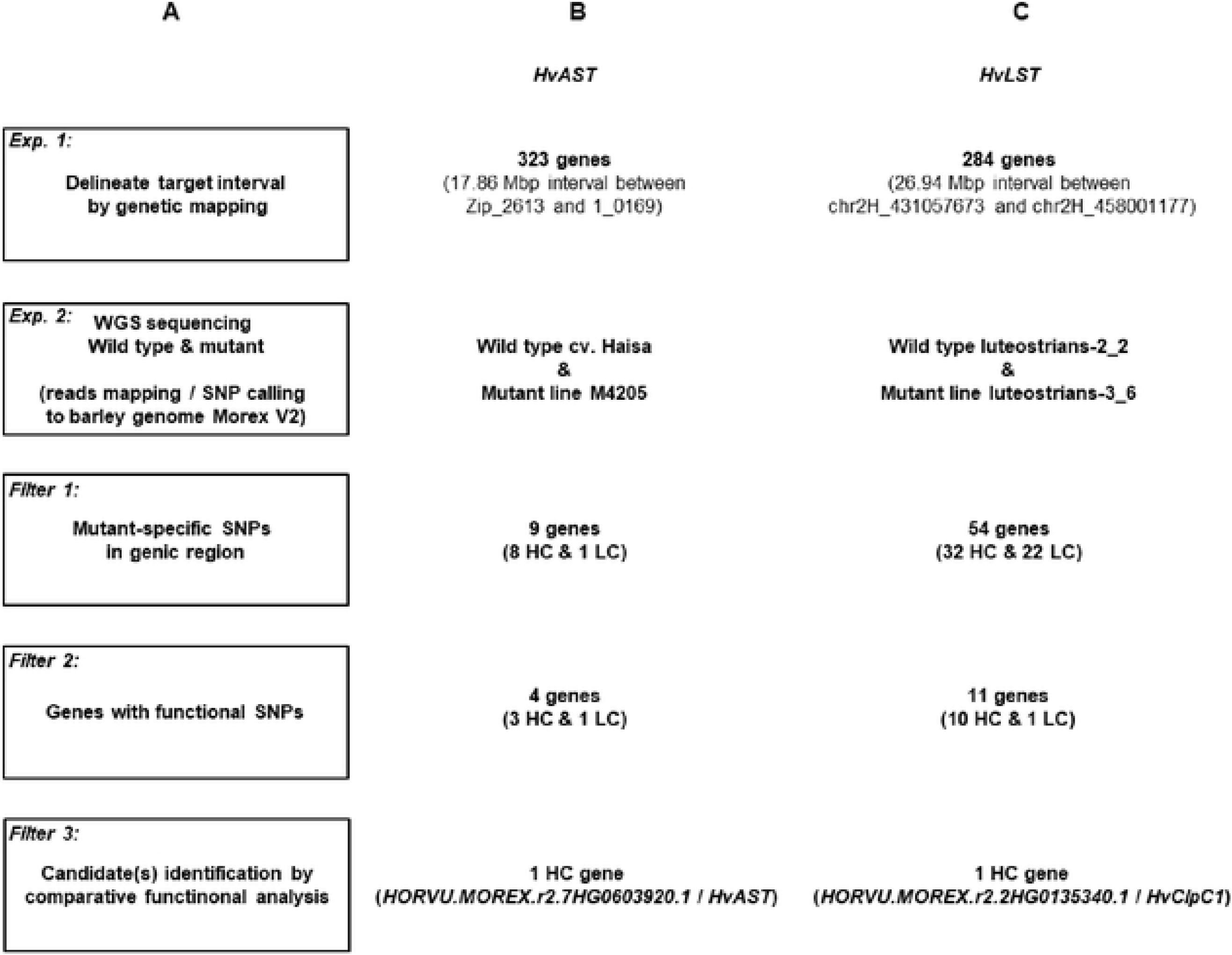
Workflow of candidate gene identification in barley photosynthetic mutants exemplified for *HvAST* and *HvLST*. (A) The initial step of the strategy built on low-resolution genetic mapping. Whole genome re-sequencing data for the mutant and its corresponding wild type genotype was then generated for mapping and variation calling against the Morex v2 reference sequence (Monat et al., 2019). (B) Candidate gene identification was exemplified on the basis of the previously cloned gene *albostrians* (*HvAST)*. 323 genes were annotated for the physical interval of ~18 Mbp initially delimited by low-resolution genetic mapping. SNP variation was found in 9 genes, while only 4 genes carried non-synonymous or other deleterious mutations. A single candidate gene (the confirmed gene *HvAST*) could be delimited based on available functional annotation information indicating a role in plastid biology / photosynthesis. (C) A similar strategy was applied to the cloning of *luteostrians*. 284 genes were annotated for the initial genetic interval. Eventually, a single candidate gene with predicted functional annotation for a role in plastid biology / photosynthesis and non-synonymous / deleterious mutation could be spotted. HC, high-confidence gene; LC, low-confidence genes. Criteria for classification of HC and LC refers to Mascher et al. (2017).

### Whole Genome Re-sequencing Identifies *HvClpC1* as *HvLST* Candidate Gene

Based on the annotated reference genome sequence of barley (Monat et al., 2019) the genetic interval of *HvLST* delimited by the closest flanking molecular markers is annotated with 284 genes (Figure 5A and 5C), a number too large for realistically spotting directly a candidate gene for *HvLST*. One option, higher resolution genetic mapping of the gene *HvLST* by screening for recombinants in a much larger mapping population was rejected due to the unfavorable genetic placement of the gene in a region of very low recombination. Instead, we evaluated the possibility to survey directly for mutations in any of the predicted genes in the *HvLST* interval by high-throughput re-sequencing.

First, we simulated the feasibility of the approach based on existing data for the variegation mutant *albostrians* (Li et al., 2019c). Previously, low-resolution mapping with 91 F2 genotypes delimited the gene *HvCMF7* to a 6.05 cM genetic interval between flanking markers Zip_2613 and 1_0169. Anchoring the flanking markers to the barley reference genome (Monat et al. 2019) revealed a 17.86 Mbp physical interval comprising 323 genes (Figure 5A-B); comparable to the situation for *HvLST*. Next, we used whole genome re-sequencing data for the mutant M4205 (original *albostrians* mutant) and barley cv. Haisa (genetic background used for induction of the *albostrians* mutant). In an initial screen we filtered M4205-specific SNPs in genic regions revealing nine candidate genes. The second step of filtering for functional SNPs (e.g., leading to non-synonymous exchange of amino acid, change of splice junction, or premature stop) retrieved four candidate genes (Figure 5A-B; Table 3 and Supplemental Table 3). Based on functional annotation of these four genes, a single gene, *HORVU.MOREX.r2.7HG0603920.1*, homolog to the Arabidopsis gene *CIA2*, was suggested as the most promising *albostrians* candidate, supported by photosynthesis-related pale green phenotype of the Arabidopsis *cia2* mutant (Sun et al., 2001). *HORVU.MOREX.r2.7HG0603920.1* in fact represents the genuine *albostrians* gene, as was verified by independent mutant analysis (Li et al., 2019c). In conclusion, based on a delimited genetic interval, survey sequencing of wild type and mutant genetic background may provide a direct path for candidate gene identification for photosynthesis related phenotype mutants in barley.

**Table 3.**
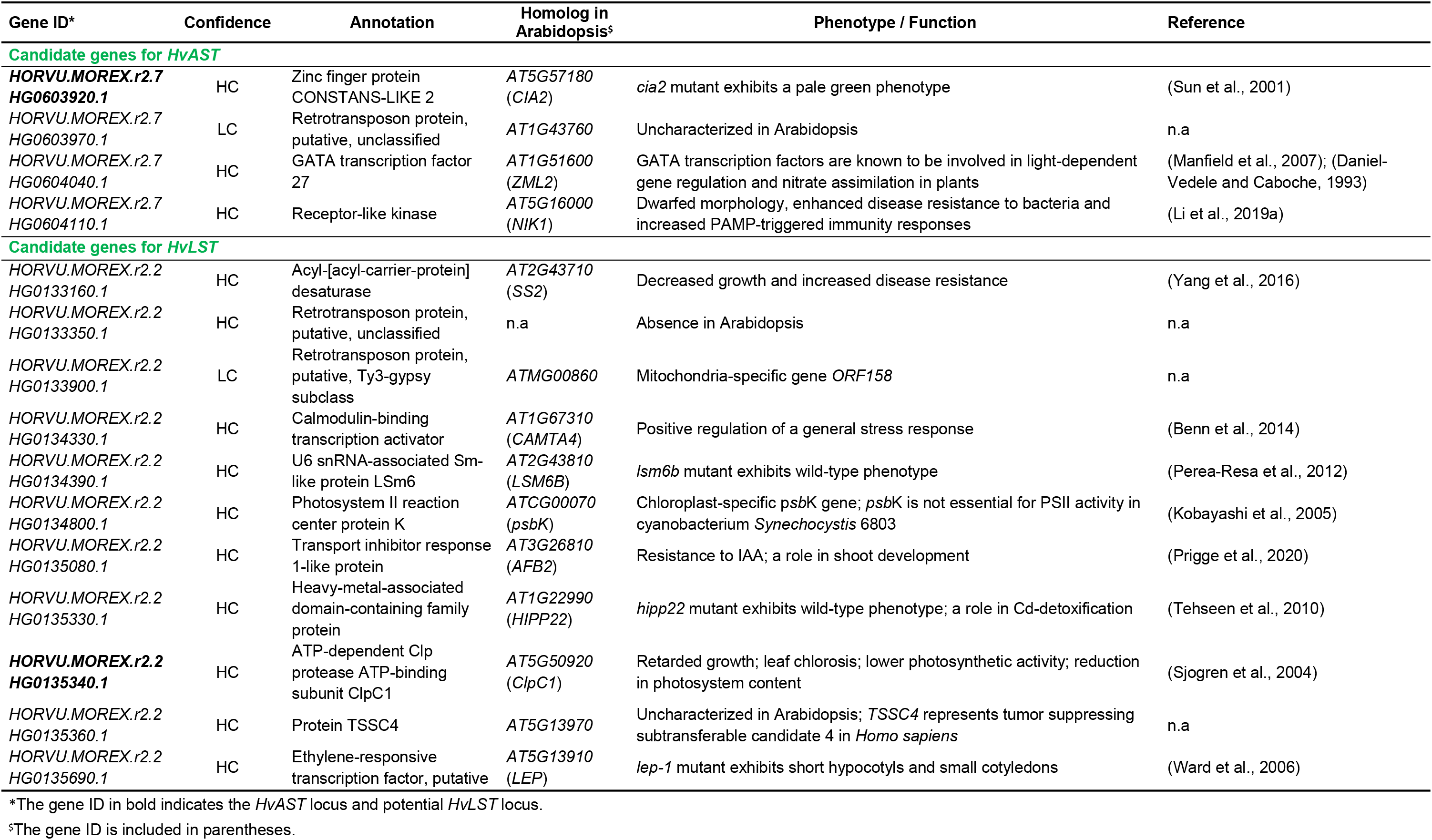
Summary of candidate genes for *HvAST* and *HvLST*.

We adopted this approach to identify a *HvLST* candidate gene. A wild type (line luteostrians-2_2, *LST*/*LST*) and a mutant genotype (line luteostrians-3_6, *lst*/*LST*) were selected from the originally segregating *luteostrians* mutant, to ensure both analyzed genotypes sharing the same genetic background of line MC20. In contrast to the previously described simulation for the gene *albostrians*, the initial screening exercise for sequence polymorphisms between mutant and wild type had to consider the fact that homozygous *lst*/*lst* is embryo lethal, hence functional polymorphisms were expected to be present at heterozygous state. Consequently, 54 out of the 284 genes, identified for the *luteostrians* mapping interval, carried heterozygous SNPs specific to the *luteostrians* mutant. For a shortlist of eleven genes, the observed SNP was predicted to induce a putative functional change (Figure 5A and 5C; Table 3 and Supplemental Table 3). BLASTp search (Mount, 2007) against the Arabidopsis proteome revealed presence of orthologs for 8 of the 11 candidate genes. None of the remaining three genes likely represented a genuine *HvLST* candidate as they either encoded for a putative retrotransposon protein in barley or showed similarity to an organelle gene lacking essential function in chloroplast biogenesis in Arabidopsis (Table 3 and Supplemental Table 3). We then inspected functional annotation information and, if available, phenotype information for mutants of the narrowed shortlist of eight Arabidopsis genes. One of these genes *AT5G50920* encodes an ATP-dependent Clp protease ATP-binding subunit ClpC1. Mutants of this gene exhibited a pale-yellow phenotype with reduced photosynthetic performance (Sjogren et al., 2004). Notably, inactivation of the *ClpP* genes (e.g., *ClpP4*, *ClpP5*, *ClpP6*) in Arabidopsis resulted in embryo lethality and antisense repression lines exhibited a variegated ‘yellow-heart’ phenotype (Clarke et al., 2005). In conclusion, *HORVU.MOREX.r2.2HG0135340.1* (*HvClpC1*), homolog (putative ortholog) of Arabidopsis *ClpC1*, represented the most likely candidate for the gene *luteostrians*. The predicted heterozygous SNP in *HvClpC1* was confirmed by Sanger sequencing (Supplemental Table 4). The *luteostrians* mutant carries a heterozygous SNP at position 2,078 (i.e., G2078A; coordinate refers to the coding sequence), consequently, changing glycine into aspartic acid at position 693 (i.e., G693D) (Figure 6B).

**Figure 6.**
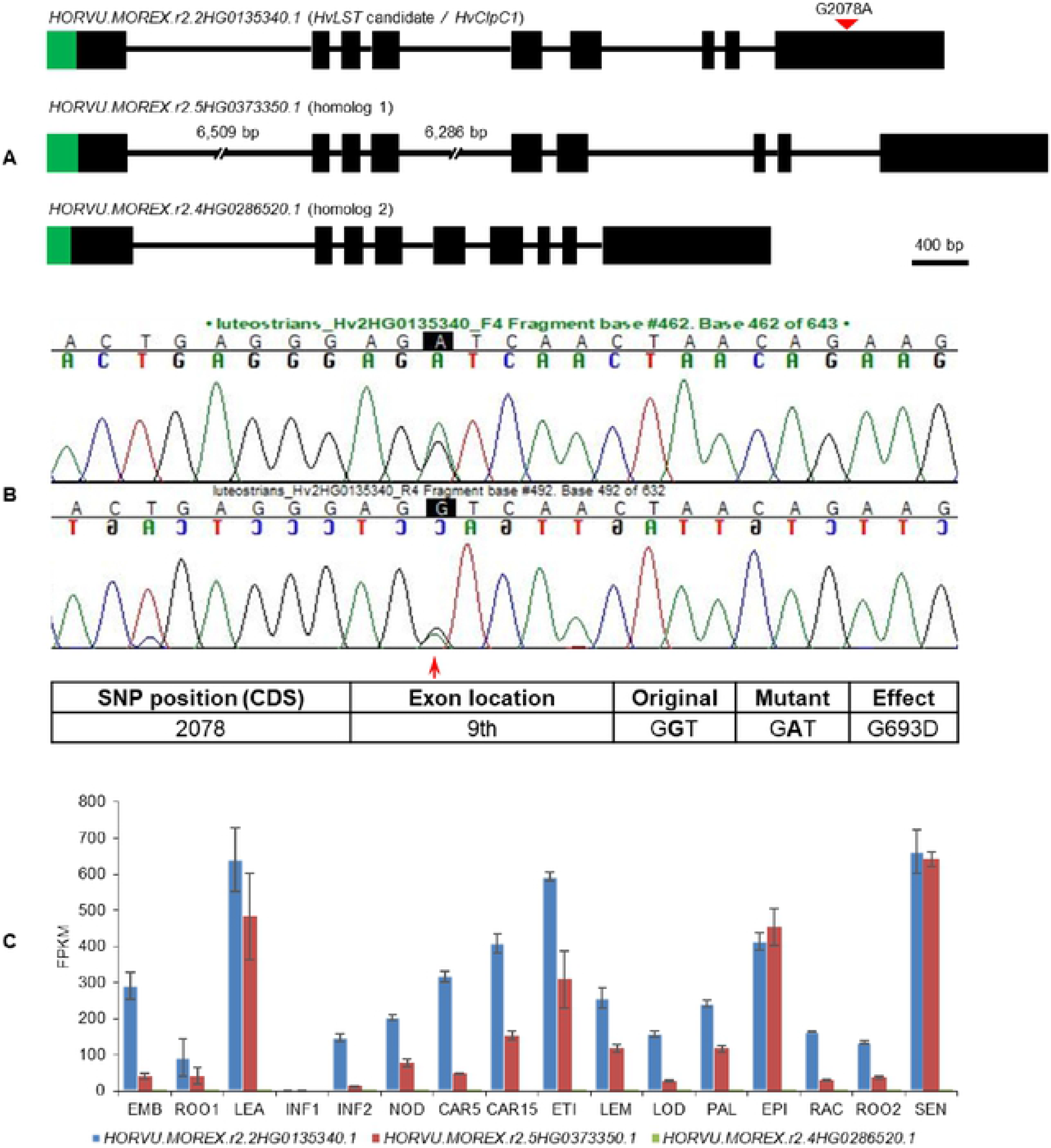
Validation of the heterozygous SNP of *HvClpC1* by Sanger sequencing and expression profiles of *HvClpC1* and homologs. (A) Gene structure of *HvLST* candidate (*HvClpC1*) and its two closest homologs. Black boxes indicate exons and horizontal lines indicate introns. Green areas indicate chloroplast transit peptides as predicted by ChloroP (Emanuelsson et al., 1999). The first and second introns of homolog 1 are not drawn at scale as indicated by the interrupted lines; the actual length is shown above gaps, respectively. Red filled triangle indicates SNP position of *HvClpC1*. (B) Chromatogram of Sanger sequencing. Red arrow indicates position of the heterozygous SNP of *HvClpC1* in the original mutant line luteostrians-1_1. Details of the SNP are illustrated in the table below. (C) Expression profiles of *HvClpC1* and its two closest homologs. The expression levels are given as fragments per kilobase of exon per million reads mapped (FPKM) across sixteen different tissues or developmental stages. The data was taken from Mascher et al. (2017). EMB, 4-day embryos; ROO1, roots from seedlings (10 cm shoot stage); LEA, shoots from seedlings (10 cm shoot stage); INF1, young developing inflorescences (5 mm); INF2, developing inflorescences (1-1.5 cm); NOD, developing tillers, 3rd internode (42 DAP); CAR5, developing grain (5 DAP); CAR15, developing grain (15 DAP); ETI, etiolated seedling, dark condition (10 DAP); LEM, inflorescences, lemma (42 DAP); LOD, inflorescences, lodicule (42 DAP); PAL, dissected inflorescences, palea (42 DAP); EPI, epidermal strips (28 DAP); RAC, inflorescences, rachis (35 DAP); ROO2, roots (28 DAP); SEN, senescing leaves (56 DAP).

The *HvLST* candidate *HvClpC1* has two close homologs in barley, *HORVU.MOREX.r2.5HG0373350.1* (homolog 1) and *HORVU.MOREX.r2.4HG0286520.1* (homolog 2), which share 85.57% and 74.26% amino acid identity with HvClpC1, respectively. All three homologs share the same gene structure with nine exons. In line with the chloroplast localization of ClpC1 in Arabidopsis (Sjogren et al., 2014), ChloroP predicts presence of a chloroplast transit peptide for all three homologs in barley (Figure 6A), potentially suggesting a chloroplast localization.

We looked up the expression profile of *HvClpC1* and its two close homologs in BARLEX database (Colmsee et al., 2015). *HvClpC1* shows ubiquitous expression in all examined tissues except young developing inflorescences (INF1) (Figure 6C). The expression pattern of homolog 1 resembles that of *HvClpC1* but in most tissues at a distinctly lower level. In contrast, expression of homolog 2 is barely detectable across all the samples.

## Discussion

Induced mutants are important tools for elucidating basic principles in the biology of photosynthesis and organellogenesis in plants. Recently, we reported the cloning and characterization of the gene which bears a mutation in the well-known green-white striped mutant *albostrians* of barley (Li et al., 2019c). Here we introduce, based on low-resolution mapping, whole genome re-sequencing and homology search to photosynthesis or chloroplast related mutants in Arabidopsis, a highly promising candidate gene *HvClpC1* for the related yellow-green variegation mutant *luteostrians*.

### Protease ATP-binding Subunit *HvClpC1* Represents the Perfect Candidate Gene for *HvLST*

Low-resolution mapping delimited *HvLST* to a large interval spanning a physical distance of 26.9 Mbp comprising a total of 284 genes. By comparative whole genome re-sequencing of wild type and mutant lines, a homolog of the Arabidopsis gene *AT5G50920* was identified as candidate *LUTEOSTRIANS* gene. *AT5G50920* belongs to the ATP-dependent protease family and encodes the ATP-binding subunit ClpC1. The *clpC1* null mutant exhibited a homogenous pale-yellow phenotype (Sjogren et al., 2004). The distinct phenotypes of *ClpC1* knockout lines in Arabidopsis and *luteostrians* barley are reminiscent of the recently reported orthologous pair of genes of the variegated *albostrians* barley mutant and its Arabidopsis homolog pale green mutant *cia2* (Sun et al., 2001; Li et al., 2019c).

The CLP protease system is a central component of the chloroplast protease network (Olinares et al., 2011). It plays essential roles in coordinating plastid proteome dynamics during developmental transitions such as embryogenesis and leaf development (Nishimura and van Wijk, 2015). Genetic studies demonstrated that members of the CLP protease family contribute to chloroplast biogenesis (Constan et al., 2004; Sjogren et al., 2004) and embryogenesis (Kovacheva et al., 2007; Kim et al., 2009). ClpC1 plays important roles on chlorophyll biosynthesis as it controls turnover of glutamyl-tRNA reductase (GluTR) and chlorophyllide *a* oxygenase (CAO); GluTR and CAO catalyse an early step of the tetrapyrrole biosynthesis pathway and the conversion of chlorophyllide *a* to *b*, respectively (Nakagawara et al., 2007; Apitz et al., 2016; Rodriguez-Concepcion et al., 2019). Although single mutants were viable, siliques of the double mutant *clpC1clpC2* (previously described as *hsp93-V / hsp93-III-2*) contained aborted seeds (because of a block in the zygote-embryo transition) and failed ovules (because of a moderate defect in female gametophytes) (Kovacheva et al., 2007). This suggested that both homologs *ClpC1* and *ClpC2* confer important functions to cell viability during gamete stage and early zygotic stage during embryogenesis. Moreover, the proteome phenotype of *clpC1clpS1* suggested a ClpS1 and ClpC1 interaction effect on plastid gene expression components and nucleoid interactors, including RNA processing and editing, as well as 70S ribosome biogenesis (Nishimura et al., 2013). So far, biochemical and genetic evidences gathered for Arabidopsis single and double mutants related to *clpC1* were in line with the observed yellow-striped, chloroplast ribosome deficient and embryonic lethal phenotype (at homozygous state) of the *luteostrians* mutant. Mutants of related genes showed also variegation in Arabidopsis, e.g., Arabidopsis mutant *yellow variegated 2 (var2*) exhibited a patched phenotype (Chen et al., 2000). *VAR2* encodes an ATP-dependent metalloprotease FtsH2. Together with the CLP and the LON family proteases, FtsH2 belongs to the ATP-dependent protease AAA+ (ATPase associated with various cellular activities) superfamily (van Wijk, 2015). FtsH2 showed functional redundancy with FtsH8 and a threshold model (i.e., level of FtsH protein complexes formed in the thylakoid membrane) was proposed for the underlying mechanism of *var2* leaf variegation (Aluru et al., 2006). In analogy to the barley mutant *albostrians* (Li et al., 2019c), green and yellow sectors of *luteostrians* have the same genotype. Furthermore, no differences could be determined for the chloroplast ultrastructure and plastid rRNA content between wild-type leaves and green sectors of striped *luteostrians* leaves. Therefore, variegation of *luteostrians* leaves may be caused by a threshold-dependent mechanism.

Though *HvClpC1* represents the most promising candidate gene for *HvLST*, further experimental evidence is required to confirm this hypothesis. As previously demonstrated, reverse genetic approaches like TILLING or site-directed mutagenesis using Cas9 are viable options in barley to further check the identity of the candidate gene with *LUTEOSTRIANS* (Gottwald et al., 2009; Lawrenson et al., 2015; Li et al., 2019c).

### Cloning-by-sequencing in Barley During the Post-NGS Era

In this study, we demonstrated that a combination of low-resolution genetic mapping and comparative whole-genome skim sequencing analysis of mutant and wild type can serve as an effective strategy for gene cloning in barley, especially for genes that share a highly conserved function in plants, including those involved in photosynthesis and chloroplast differentiation and development. The probability of having available functional characterization data for such genes in model plants is high and there is increased probability for seeing related or conserved phenotypic effects in both model and non-model plants. By adopting this strategy, we pinpointed the *albostrians* gene *HvCMF7* among 323 candidates within a 17.86 Mbp interval delimited with 91 F2 genotypes, circumventing the time-consuming and laborious fine-mapping process as performed previously (Li et al., 2019c). Implementation of the pipeline in case of the gene *luteostrians* suggested *HvClpC1* to be the likely causative gene underlying the yellow-stripe variegation phenotype. Notably, the cloning strategy is independent of the effect of genomic location. Compared to the barley genome-wide average ratio of physical to genetic distance of 4.4 Mb/cM (Kunzel et al., 2000; Mascher et al., 2017), *HvAST* and *HvLST* are located in genomic regions with either increased (2.95 Mb/cM) or suppressed (36.35 Mb/cM) rates of recombination, respectively. The success of candidate gene identification in both cases can be attributed to the achieved high-quality and completeness of the reference genome of barley (Mascher et al., 2017; Monat et al., 2019). Whole genome re-sequencing of wild type and mutant parental lines revealed four and eleven candidates for *HvAST* and *HvLST*, respectively. To further narrow down the shortlist to a single candidate gene, it was critical that functional analyses for the putative Arabidopsis homologs were available. In both cases, the best candidate gene showed a chlorophyll-deficient phenotype for mutants of the respective Arabidopsis homolog.

Genetic mapping information is available for 881 barley mutants of the so-called Bowman near-isogenic lines (NILs) (Druka et al., 2011); among them, 142 mutants with pigmentation phenotype. Although the original genetic background of the Bowman NILs is represented by over 300 different barley genotypes, approximately one-half of the mutants (426/881) were identified in one of eight barley cultivars (i.e., Bonus, Foma, Betzes, Akashinriki, Morex, Steptoe, Volla and Birgitta) (Druka et al., 2011). Since leveraging of the high-quality barley reference genome sequence (Mascher et al., 2017; Monat et al., 2019), whole-genome resequencing of the parental lines would help to filter background mutations and facilitate candidate gene identification and isolation. Therefore, it is conceivable that gene identification in barley, given this pilot attempt based on chlorophyll / photosynthesis related mutants, reached to a point to become even faster and more systematic now.

## Materials and Methods

### Plant Materials and Growth Conditions

The *luteostrians* mutant was derived from line MC20 [Institute of Genetics ‘Ewald A. Favret’ (IGEAF), INTA CICVyA, mutant accession number] by chemical mutagenesis using sodium azide. MC20 was derived from the spring barley cv. Maltería Heda by gamma ray mutagenesis. Two mapping populations were constructed. The six-rowed spring barley (*Hordeum vulgare*) cv. ‘Morex’ and two-rowed spring barley cv. ‘Barke’ were used as maternal parent in crossings with the variegated mutant line either *luteostrians*-1 (*lst/LST*) or *luteostrians*-3 (*lst/LST*), respectively. The derived F1 progeny were self-pollinated to generate F2 mapping populations designated as ‘ML’ (Morex x *luteostrians*) and ‘BL’ (Barke x *luteostrians*). All the F2 and F3 individuals were grown under the greenhouse condition with a day/night temperature and photoperiod cycle of 20°C/15°C and 16-h light/8-h darkness, respectively. Supplemental light (at 300 μmol photons m^−2^ s^−1^) was used to extend the natural light with incandescent lamps (SON-T Agro 400, MASSIVEGROW).

For ribosomal RNA analysis, wild type (*luteostrians*-2; genotype *LST*/*LST*) and mutant (*luteostrians*-1; genotype *lst*/*LST*) lines were grown under the greenhouse light condition. For dark treatment, seeds were germinated within a carton box wrapped with aluminum foil under the greenhouse condition mentioned above.

### Phenotyping

The trait ‘leaf color’ was scored for the F2 mapping populations at first, second and third leaf seedling stages. The phenotype of the seedlings was classified into two categories: green and variegated (i.e., green-yellow striped). Green phenotype defined all the three leaves of each seedling were purely green; and variegated phenotype defined green-white striped pattern can be observed on all or any of the three leaves. According to inheritance pattern of the causal gene *luteostrians* (Figure 1B), variegated plants are heterozygous for the mutant allele (*lst/LST*). Green plants, however, can be either wild type (*LST/LST*) or heterozygous (*LST/lst*). Therefore, the *luteostrians* genotype of all the F2 plants was determined by phenotyping 32 seedlings of each corresponding F3 family. A wild type (*LST/LST*) F2 would produce 100% green progeny, progeny of heterozygous (*LST/lst*) F2 green plant would follow a Mendelian segregation (green:variegated = 2:1).

### Ribosomal RNA Analysis

Total RNA was extracted using TRIzol reagent (Invitrogen, Braunschweig, Germany) following manufacturer’s instructions. Initially, concentration of the RNA was determined by help of a Qubit^®^ 2.0 Fluorometer (Life Technologies, Darmstadt, Germany) according to manufacturer’s instructions. RNA samples were further diluted within a quantitative range of 1 - 10 ng/μL. RNA quality and quantity were then measured using Agilent High Sensitivity RNA ScreenTape following the manufacturer’s manual (Agilent, Santa Clara, USA).

### Ultrastructural analysis

Primary leaves were collected from 7-day-old seedlings of wild type and variegated mutant (*luteostrians*-1). Preparation of 1-2 mm^2^ leaf cuttings for ultrastructure analysis including aldehyde/osmium tetroxide fixation, dehydration, resin infiltration and ultramicrotomy were performed as previously described (Li et al., 2019b).

### DNA Isolation

Genomic DNA was extracted from primary leaves of 7-day-old seedlings using a GTC-NaCL method in 96-well plate format. Frozen leaf samples were homogeneously crushed by help of a Mixer Mill MM400 (Retsch GmbH, Haan, Germany) for 30 s at 30 Hz. Add 600 μL of preheated (65°C for 1 hour) GTC extraction buffer (1 M guanidine thiocyanate, 2 M NaCl, 30 mM NaCOOH pH = 6.0, 0.2% Tween 20) to the frozen powder and mixed thoroughly for 30 s at 30 Hz under Mixer Mill MM400. Centrifugation at 2500 x *g* for 10 min at 4°C after incubation the samples at 65°C for 1 hour. Transfer 480 μL of supernatant to a 96-well EconoSpin plate (Epoch Life Science, Texas, USA). Vacuum the EconoSpin plate on the vacuum manifold (MACHEREY-NAGEL, Dueren, Germany) until no droplet drop down. Wash the DNA samples twice with 880 μL washing buffer (50 mM NaCl, 10 mM Tris-HCl pH = 8.0, 1 mM EDTA, 70% ethanol). Place the EconoSpin plate onto a 96-well Microtiter™ microplate (Thermo Fisher Scientific, Braunschweig, Germany) and centrifugation at 2500 x *g* for 3 min to remove residual wash solution. Dissolve DNA with 100 μL preheated (65°C for 1 hour) TElight buffer (0.1 mM EDTA, 10 mM Tris-HCl pH = 8.0). The isolated DNA were used for downstream analysis or put at −20°C for long term storage.

### Genotyping-by-sequencing

For the genetic mapping of *HvLST*, GBS were prepared from genomic DNA extracted as described previously, digested with *Pst*I and *Msp*I (New England Biolabs, Frankfurt am Main, Germany) following published procedure (Wendler et al., 2015). DNA samples were pooled in an equimolar manner per lane and sequenced on Illumina HiSeq2500 for 107 cycles, single read, using a custom sequencing primer. The GBS reads were aligned to the reference genome of barley as described by Milner et al. (2019). Reads were trimmed using cutadapt (Martin, 2011), mapped with BWA-MEM version 0.7.12a (Li and Durbin, 2009) to barley reference genome (Monat et al., 2019) and the resulting bam files were sorted using Novosort (Novocraft Technologies Sdn Bhd, Selangor, Malaysia). Variants were called with samtools version 1.7 and bcftools version 1.6 (Li, 2011) and filtered following the protocol of Milner et al. (2019) for a minimum depth of sequencing of six to accept a genotype call, a maximum fraction of heterozygous call of 60% and a maximum fraction of 20% of missing data. In the case of ‘BL’ population, SNP calls were converted to reflect the polymorphisms between the two parents, using the calls for barley cv. Barke.

### Genetic Linkage Map Construction

Genetic linkage groups were constructed by help of the JoinMap^®^ 4.1 software (Van Ooijen, 2006) following the instruction manual. Homozygous wild type and heterozygous allele calls were defined as A and H, respectively; missing data was indicated by a dash. A permissive threshold of 5% missing data for both molecular marker and F2 genotype was applied to the datasets. Regression mapping algorithm and Kosambi’s mapping function were chosen for building the linkage maps. Markers were grouped into seven groups based on Logarithm of Odds (LOD ≥ 6) groupings. The seven linkage groups were in corresponding to the barley chromosomes according to the locus coordinates determined during read mapping to the reference genome of barley (Monat et al., 2019). Visualization of maps derived from JoinMap^®^ 4.1 was achieved by MapChart software (Voorrips, 2002).

### KASP Assay

Sequence 50 bp upstream and downstream of the SNP was extracted from the barley reference genome (Monat et al., 2019). Allele-specific forward primers and one common reverse primer were designed by help of the free assay design service of 3CR Bioscience (https://3crbio.com/free-assay-design/) (Supplemental Table 4). KASP primer mix was prepared in a volume of 100 μL containing 12 μM of each allele-specific forward primer and 30 μM common reverse primer. KASP-PCR reactions were performed in a total volume of 5 μL containing 2.5 μL 2X PACE-IR™ Genotyping Master Mix (001-0010, 3CR Bioscience, Essex, UK), 0.07 μL KASP primer mix, and 40 ng of template DNA. PCR program was used with a HydroCycler (LGC, Teddington, UK): initial denaturation at 94°C for 15 min followed by 10 cycles at 94°C for 20 s, at 65 to 57°C (−0.8°C/cycle) for 1 min, and proceeded for 30 cycles at 94°C for 20 s, at 57°C for 1 min, and followed at 30°C for 30 s. In case of the genotyping clusters not well separated, an additional PCR was performed with 6 cycles at 94°C for 20 s, at 57°C for 1 min. Pre-read and post-read of the fluorescence signals were performed on ABI 7900HT instrument (Applied Biosystems, Thermo Fisher Scientific).

### Whole-genome Shotgun Sequencing and Data Analysis

DNA was isolated according to the protocol of Doyle and Doyle (1990) from leaf samples collected from one-week-old, greenhouse-grown seedlings of barley cv. Haisa, *albostrians* mutant M4205, and lines luteostrians-2_2 and luteostrians-3_6. For lines luteostrians-2_2 and luteostrians-3_6, DNA was fragmented (200 - 500 bp) with ultrasounds, then 150 bp paired-end libraries were prepared according to Illumina standard protocol and sequenced with Illumina Hiseq X Ten. For Haisa and *albostrians* mutant M4205, 350 bp paired-end libraries were prepared and sequenced according to the protocol as described previously (International Barley Genome Sequencing Consortium, 2012). Reads mapping and variants calling to the barley reference genome were performed as above described for GBS data. Notably, re-sequencing data of barley cv. Barke, maternal parent of the BL mapping population (for *luteostrians*) and BM mapping population (for *albostrians*) (Li et al., 2019c) was also included for SNP calling for identification of mutant-specific SNPs. Functional effect of the mutant-specific SNPs was annotated by help of SNPeff version 4.3 (Cingolani et al., 2012).

## Acknowledgements

We gratefully acknowledge Mary Ziems and Heike Harms for their technical support in maintaining plant material, performing crossings for construction of the mapping populations; Manuela Kretschmann for assistance in KASP genotyping; Claudia Riemey for help on electron microscopy; Heike Mueller for photography. The work was supported by a fellowship of the China Agriculture Research System (CARS-05) and the Agricultural Science and Technology Innovation Program of CAAS to G.G. and by the Deutsche Forschungsgemeinschaft grant (DFG) 1102/13-1 to N.S.

## Author Contributions

M.L. and N.S. conceived the research; A.R.P. performed initial genetic characterization of the *luteostrians* mutant; M.L., G.G. and M.M. performed experiments; M.L., H.P. and G.G. analyzed data; M.L. wrote the manuscript with contributions of H.P.; M.L., T.B. and N.S. revised the manuscript.

## Supplemental Data

**Supplemental Figure 1.** Genetic mapping of the *HvLST* gene.

**Supplemental Table 1.** Summary of SNP markers derived from genotyping-by-sequencing.

**Supplemental Table 2.** Graphical genotype of selected F2 recombinants.

**Supplemental Table 3.** Summary of candidate genes for *HvAST* and *HvLST*.

**Supplemental Table 4.** Primers used in this study.

